# Effect of 5-*trans* isomer of arachidonic acid on model liposomal membranes studied by a combined simulation and experimental approach

**DOI:** 10.1101/279422

**Authors:** Ioanna Tremi, Dimitrios Anagnostopoulos, Ellas Spyratou, Paraskevi Gkeka, Alexandros G. Georgakilas, Chryssostomos Chatgilialoglu, Zoe Cournia

## Abstract

Unsaturated fatty acids are found in humans predominantly in the *cis* configuration. Fatty acids in the *trans* configuration are primarily the result of human processing (*trans* fats), but can also be formed endogenously by radical stress. The *cis-trans* isomerization of fatty acids by free radicals could be connected to several pathologies. *Trans* fats have been linked to an increased risk of coronary artery disease; however, the reasons for the resulting pathogenesis remain unclear. Here, we investigate the effect of a mono *trans* isomer of arachidonic acid (C20:4-5*trans*,8*cis*,11*cis*,14*cis*) produced by free radicals in physiological concentration on a model erythrocyte membrane using a combined experimental and theoretical approach. Molecular Dynamics (MD) simulations of two model lipid bilayers containing arachidonic acid and its 5-*trans* isomer in 3% mol. were carried out for this purpose. The 5-*trans* isomer formation in the phospholipids was catalyzed by HOCH_2_CH_2_S• radicals, generated from the corresponding thiol by γ-irradiation, in multilamellar vesicles (MLVs) of SAPC. Large unilamellar vesicles were made by the extrusion method (LUVET) as a biomimetic model for *cis*-*trans* isomerization. Atomic Force Microscopy and Dynamic Light Scattering were used to measure the average size, morphology, and the z-potential of the liposomes. Both results from MD simulations and experiments are in agreement and indicate that the two model membranes display different physicochemical properties in that the bilayers containing the *trans* fatty acids were more ordered and more rigid than those containing solely the *cis* arachidonic acid. Correspondingly, the average size of the liposomes containing *trans* isomers was smaller than the ones without.

## Introduction

Unsaturated fatty acids are fundamental components of plasma cell membranes and play a crucial role in cellular functions (Ibarguren, López et al. 2014). The natural occurring geometry of fatty acids double bonds is the *cis* configuration, which can be found in most bacterial and mammalian cells. *Trans* isomers found in mammalian membranes are attributed to exogenous sources such as food intake (Judd and Clevidence 1993, Mendonça, Araújo et al. 2017). *Trans* fatty acids can enter the food market by three main sources: (i) partial hydrogenation of fats, (ii) high-temperature processing of edible oils (deodorization processes), and (iii) use of ruminant meats and dairy products that have a *trans* content due to the biohydrogenation process taking place during animal metabolism (Chatgilialoglu, Ferreri et al. 2014). Examples of mono-and polyunsaturated fatty acid (MUFA and PUFA) structures are shown in Fig. 1A with their common names.

**Fig. 1.**
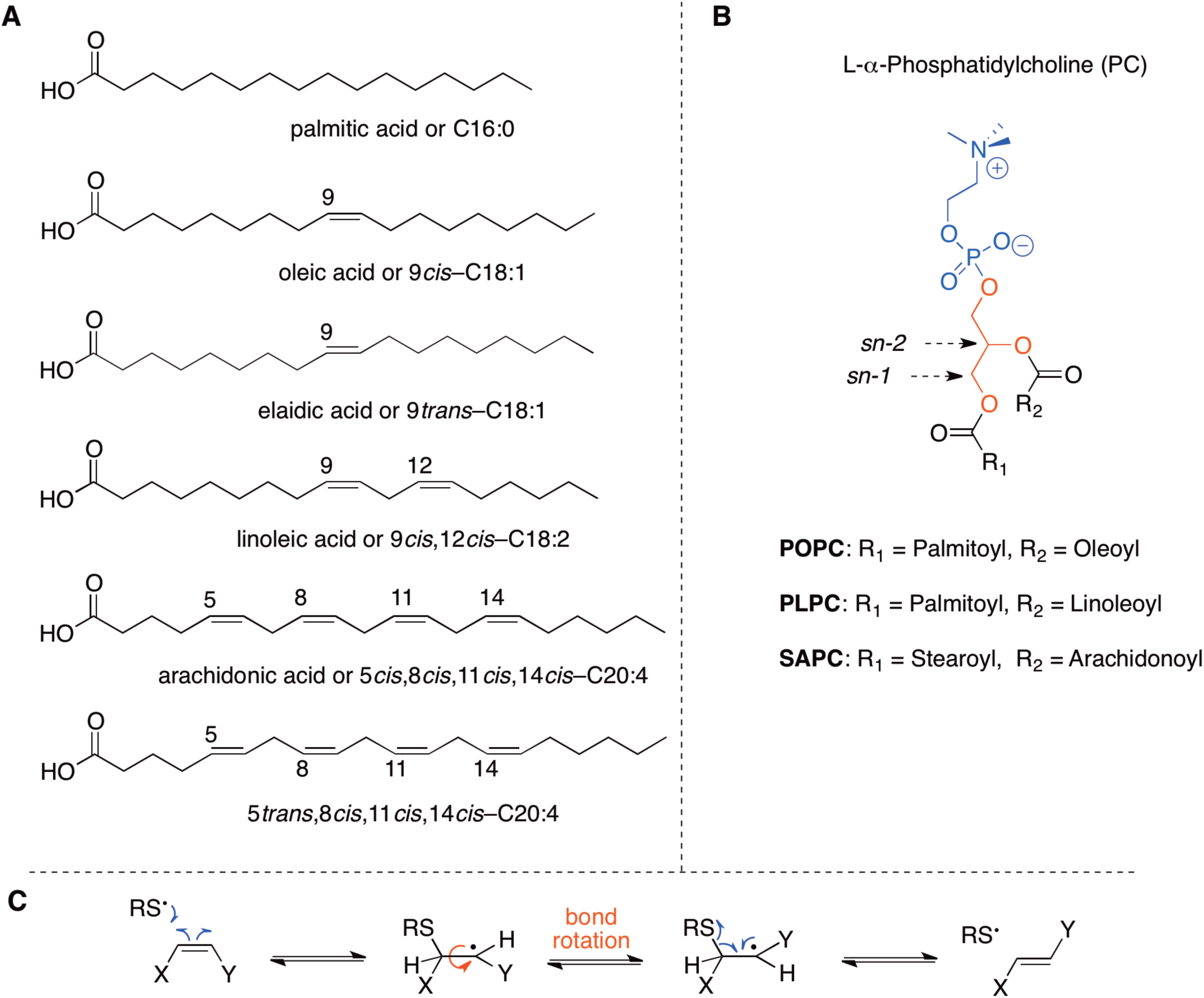
(A) Examples of natural fatty acids and some trans isomers, the abbreviations describing the position and geometry of the double bonds (e.g., 9*cis* or 9*trans*), as well as the notation of the carbon chain length and total number of unsaturations (e.g., C18:1); (B) Molecular structures of L-α-phosphatidylcholine, in the *sn-1* and *sn-2* positions of the glycerol moiety the two fatty acid residues are attached, whereas in the *sn*-3 position the polar head group is connected; (C) Reaction mechanism for the *cis-trans* isomerization catalyzed by thiyl radicals

Thiyl radical catalyzed *cis-trans* isomerization of unsaturated lipids under biomimetic conditions has been a subject of interest from some of us for the past decade (Chatgilialoglu and Ferreri 2005, Chatgilialoglu, Ferreri et al. 2014). Strong indications were provided for an endogenous origin of *trans* unsaturated fatty acids in cells (Ferreri, Kratzsch et al. 2005, Cort, Ozben et al. 2016), animals (Zambonin, Ferreri et al. 2006), and humans (Puca, Andrew et al. 2008, Ferreri and Chatgilialoglu 2012, Sansone, Melchiorre et al. 2013, Sansone, Tolika et al. 2016), based on mono-*trans* PUFA isomers as biomarkers (Ferreri, Faraone Mennella et al. 2002, Ferreri, Samadi et al. 2004, Melchiorre, Torreggiani et al. 2011). Thiyl radical catalyzed *cis-trans* isomerization of a lipid double bond is shown in Fig. 1C.

Although a relationship between *trans* fatty acids and some diseases has been established, the mechanism by which *trans* fats cause the associated pathogeneses remains elusive. It has been shown that when dietary *trans* fatty acids are incorporated into cell membranes they may lead to alteration of membrane physicochemical properties such as reduction of fluidity and permeability and formation of a more rigid packing in the lipid bilayer, which may also contribute to the observed pathogenesis (Egwim and Sgoutas 1971, Emken, Adlof et al. 1983, Wolff and Entressangles 1994). The *trans* lipid geometry resembles that of saturated lipids (Wolff and Entressangles 1994), which may mimic a decrease in the degree of unsaturation in the cell membrane that leads to damages in the cell adaptation responses (Fig. S1) (Murakami, Tsuyama et al. 2000). Although previous biophysical studies have indicated that the effect of *trans* lipids on model membranes, the molecular description of this effect is still lacking. Molecular Dynamics (MD) simulations of lipid membranes can provide atomic-level detailed description of underlying physicochemical phenomena such as lipid packing and molecular interactions. It should be noted that there is only one simulation study that investigates the effect of *trans* lipids in model membrane (Kulig, Pasenkiewicz-Gierula et al. 2016).

Several saturated and unsaturated membrane lipids are present in cell membranes and their ratios differ between different cell types (Lauritzen, Hansen et al. 2001). In this study, we focus on the erythrocyte membrane as a model system to study the effect of *trans* fats in the physicochemical properties of the cell membrane, because they have been extensively studied due to their easier biochemical analysis compared to other cells, as they lack a nucleus (Gorter and Grendel 1925, Neil W Blackstone 2007). The erythrocyte lipid composition differs between the inner and outer layer of the cell membrane; here we focus on the outer layer, in which phosphatidylcholines are mostly present (Fig. 1B) (van Meer, Voelker et al. 2008). Studies of the erythrocyte phospholipids composition shows that the distribution of fatty acids in healthy individuals is the following: saturated fatty acids (SFA) 30-45% mol., monounsaturated fatty acids (MUFA) 13-23% mol. and polyunsaturated fatty acids (PUFA) 28-39% mol. (Puca, Andrew et al. 2007, Ferreri and Chatgilialoglu 2012).

Arachidonic acid is abundant in erythrocytes, and is involved in the regulation of signaling enzymes as a key inflammatory intermediate, and can also act as a vasodilator (Ricciotti and FitzGerald 2011, Norris and Carr 2013). Arachidonic acid is also involved in metabolic processes, which also play an important role in carcinogenesis (Yarla, Bishayee et al. 2016). The oxidation of arachidonic acid and the generation of *trans* isomers may affect some of the above functions. Indeed, its four double bonds render it more susceptible to oxidation (Gardner 1989) and its 5-*trans* and 8-*trans* isomers are related to the endogenous radical-based *cis-trans* isomerization (Folch, Lees et al. 1957, Ferreri, Samadi et al. 2004). Taking into account also the close relationship between oxidative stress and lipid peroxidation with various pathological conditions including cancer, cardiovascular disease, and diabetes (Birben, Sahiner et al. 2012, Siegel, Ermilov et al. 2014), investigating the role of *cis-trans* arachidonic acid isomerization on the membrane physicochemical properties could provide additional useful information on the role of stress root conditions in health and disease.

In this paper, we have investigated, for the first time, the effect of 5-*trans* isomer of arachidonic acid (5*trans*,8*cis*,11*cis*,14*cis*-C20:4) on the physicochemical properties of model membranes. We employ a combination of theoretical and experimental approaches such as MD simulations, Atomic Force Microscopy (AFM) and Dynamic Light Scattering (DLS) experiments. It should be noted that the erythrocyte membrane model used for cis/trans effect contains 3% mol of the 5-trans isomer of arachidonic acid in the total fatty acid content of the phospholipids, which is 8-10 times more than found in healthy red blood count (Ferreri and Chatgilialoglu 2012). In summary, we report that *trans* lipids become more ordered than their corresponding *cis* form suggesting similarities between saturated lipids and *trans* fatty acids, posing thus an overall ordering effect in the model membrane. The liposomes containing the *trans* isomers are more rigid and their diameter is smaller than the ones without, in agreement with recent literature (Giacometti, Marini et al. 2017). Both AFM and DLS results are in agreement with the theoretical predictions.

## Materials and Methods

### Materials

HOCH_2_CH_2_SH and 1-palmitoyl-2-oleyl-*sn*-glycero-3-phosphocholine (POPC) were purchased from Aldrich. 1-Palmitoyl-2-linoleoyl-*sn*-glycero-3-phosphocholine (PLPC) and 1-stearoyl-2-arachidonoyl-*sn*-glycero-3-phosphocholine (SAPC) were purchased from Avanti Polar Lipids (Alabaster, AL) (see Fig 1B). All solvents were HPLC-grade and the utilized water was glass-distilled. Radiolysis was performed at room temperature (22 ± 2 °C) on 1 mL samples using a ^60^Co Gammacell. For the construction of 100 nm liposomes the Mini Extruder was used from Avanti Polar Lipids. The composition of lipids during the experiments was monitored by gas chromatography using the GC 7890? from Agilent equipped with a flame ionization detector. The column used was the DB-23 capillary column (60m × 0.25mm). The heating was carried out at a temperature of 165 °C for 3 minutes followed by an increase of 1 °C/min up to 195 °C. The temperature remained at 195 °C for 40 minutes and then it was followed by an increase of 10 °C /min up to 240 °C for 10 minutes.

### Liposome Preparation

Multilamellar vesicles of 10 mM were prepared using the lipid film hydration method following System 1 and System 2 (Table 1) (Samad, Sultana et al. 2007). A mixture of POPC (40% mol.), PLPC (30% mol.) and SAPC (30% mol.) were dissolved in a small amount of methanol:chloroform 1:2 (v/v). The solution was placed in a rotary evaporator at 40°C until a thin film is created on the wall of a flask. The resulting lipid was kept under vacuum in order to eliminate any traces of organic solvents. The lipid film was then hydrated with 1mL distilled water by mixing in a vortexing machine for 5 minutes. Then, large unilamellar vesicles were made by the extrusion method (LUVET), where the original multilamellar vesicles (MLVs) went through polycarbonate membrane filters (100 nm pore size Millipore filter). 19 passages were performed to avoid contamination of the sample by large vesicles, which might not have passed through the filter.

**Table 1:** Lipid Composition of the two studied systems

| Phospholipid (PC) | System 1 | System 2 |
| --- | --- | --- |
| POPC | 40% mol. | 40% mol. |
| PLPC | 30% mol. | 30% mol. |
| SAPC | 30% mol. | 27% mol. |
| TSAP | 0% mol. | 3% mol. |

### Synthesis of TSAP

MLVs of 10mM consisting only of SAPC lipids were prepared following the above protocol. The MLVs were diluted in the presence of 2-mercaptoethanol (HOCH_2_CH_2_SH) to reach 1mM final concentration in 2:1 ratio (Chatgilialoglu, Ferreri et al. 2014). Then, liposomes were irradiated using γ-irradiation (5 Gy/min) for 25 minutes in order to acquire the desired *cis*-*trans* percentage distribution. The isomerization was catalyzed by the corresponding HOCH_2_CH_2_S• radical (Ferreri, Costantino et al. 2001). Throughout all the experiments the composition of the liposomes as well as the generation of 5-*trans* arachidonic acid (C20:4-5*trans*, 8*cis*, 11*cis*, 14*cis*) from the irradiated SAPC liposomes is monitored by gas chromatography. A representative chromatograph of the irradiated SAPC liposomes can be seen in Fig. S2. To the phosphatidylcholine containing the 5-*trans* isomer of arachidonic acid was given the acronym TSAP.

### Liposome characterization by Atomic Force Microscopy (AFM)

The AFM observations (morphology and average size diameter) were performed with the Innova^TM^ Scanning Probe Microscope. Muscovite mica was used as a solid substrate. For each measurement, 10ul of liposome suspension was placed on the mica substrate and then the liposomes were absorbed by the hydrophilic mica. Results are presented as mean ± SD.

### Liposome Characterization by Dynamic light Scattering (DLS)

Malvern’s Zetasizer nano ZS was used to calculate the average size of the liposomes and the zeta potential in comparison with the AFM measurements. The device determines the particle size by measuring the scattering of the laser beam in the suspended particles (due to the Brownian motion of the particles). The liposomal differences are measured and the mean diameters and particle size distributions are calculated. For the measurement of the liposomes, dilution was carried out with purified water, by reaching a final concentration of 0.05 mM. Further filtration through polycarbonate filters, with a 0.22 µm pore size Millipore filter, was performed to remove particles. Results are presented as mean ± SD.

### Simulated Model and Protocol

POPC (16:0/18:1-9*cis*), PLPC (16:0/18:2-9*cis*, 12*cis*), SAPC (18:0/20:4-5*cis*, 8*cis*, 11*cis*, 14*cis*), and TSAP (18:0/20:4-5*trans*, 8*cis*, 11*cis*, 14*cis*) phospholipids were used in this study to provide a realistic composition of the erythrocyte membrane (for a list of the abbreviations, please check the SI). As TSAP we denote the SAPC phospholipids that contain the 5-*trans* isomer of arachidonic acid in the *sn*2 chain of SAPC (see Fig. 2). The composition of these model membranes was the same both for the liposomes used in the experiments and the simulated lipid bilayers, and can be seen in Table 1.

**Fig. 2.**
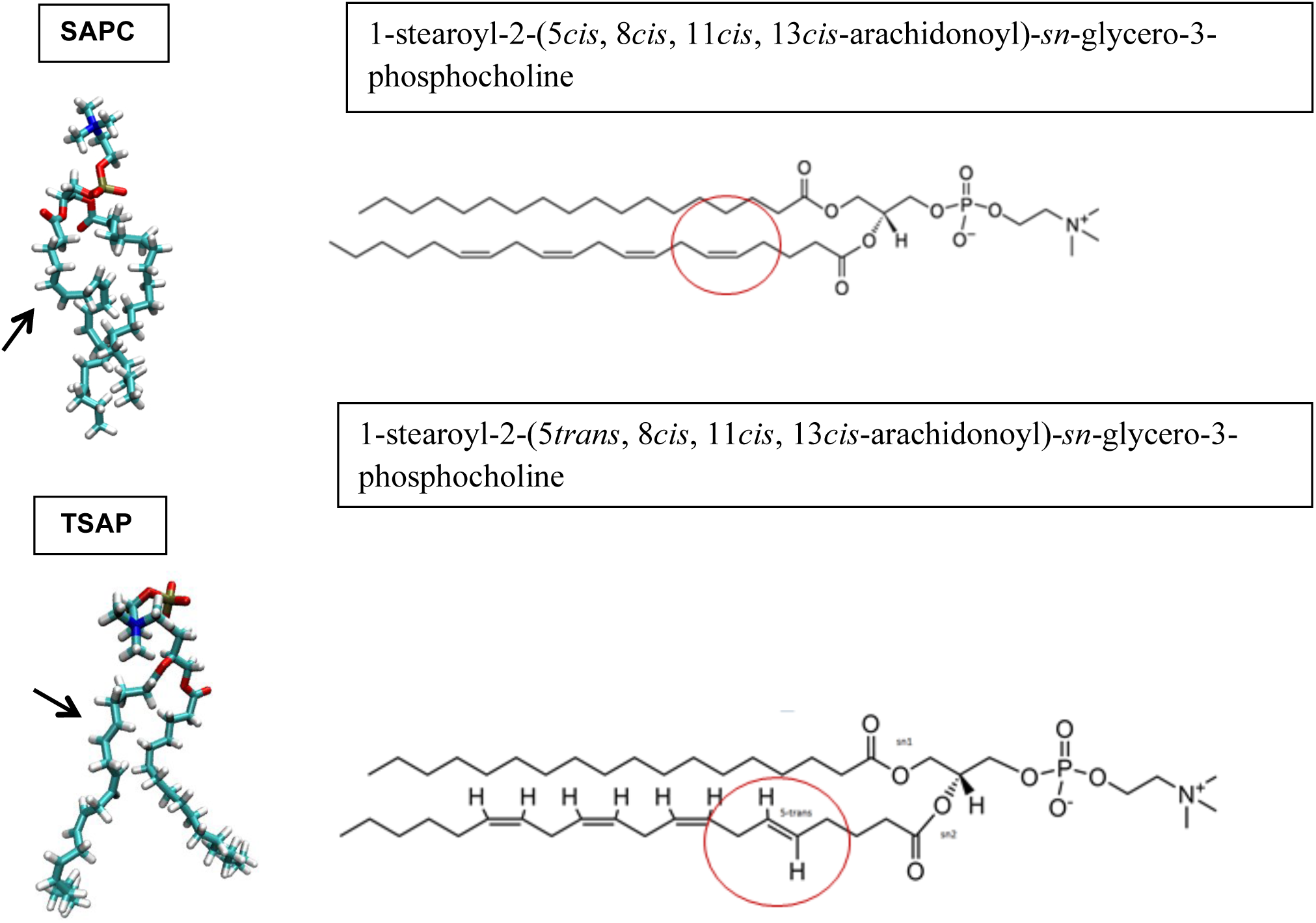
Representation of the *cis*-*trans* configuration on the *sn*2 chain of the arachidonic acid. The arrows show the isomerized double bond. Circles indicate the *cis* and *trans* isomers of the arachidonic acid on the 5 position of the *sn*2 chain.

Initially, a lipid bilayer (System 1) was generated using the charmm-gui membrane builder (Wu, Cheng et al. 2014). The membrane consisted of 520 PC (phosphatidylcholine) molecules (208 POPC, 156 PLPC, and 156 SAPC) and a snapshot can be seen in Fig. 3. In our initial system, all the double bonds of the fatty acids were in the *cis* configuration. In order to create the *trans* configuration SAPC dihedral restraints were applied to restraint the first double bond to 180^°^ (Fig. 2) (Murzyn, Rog et al. 2001). For the procedure we followed to create the *trans* configuration, see the SI.

**Fig. 3.**
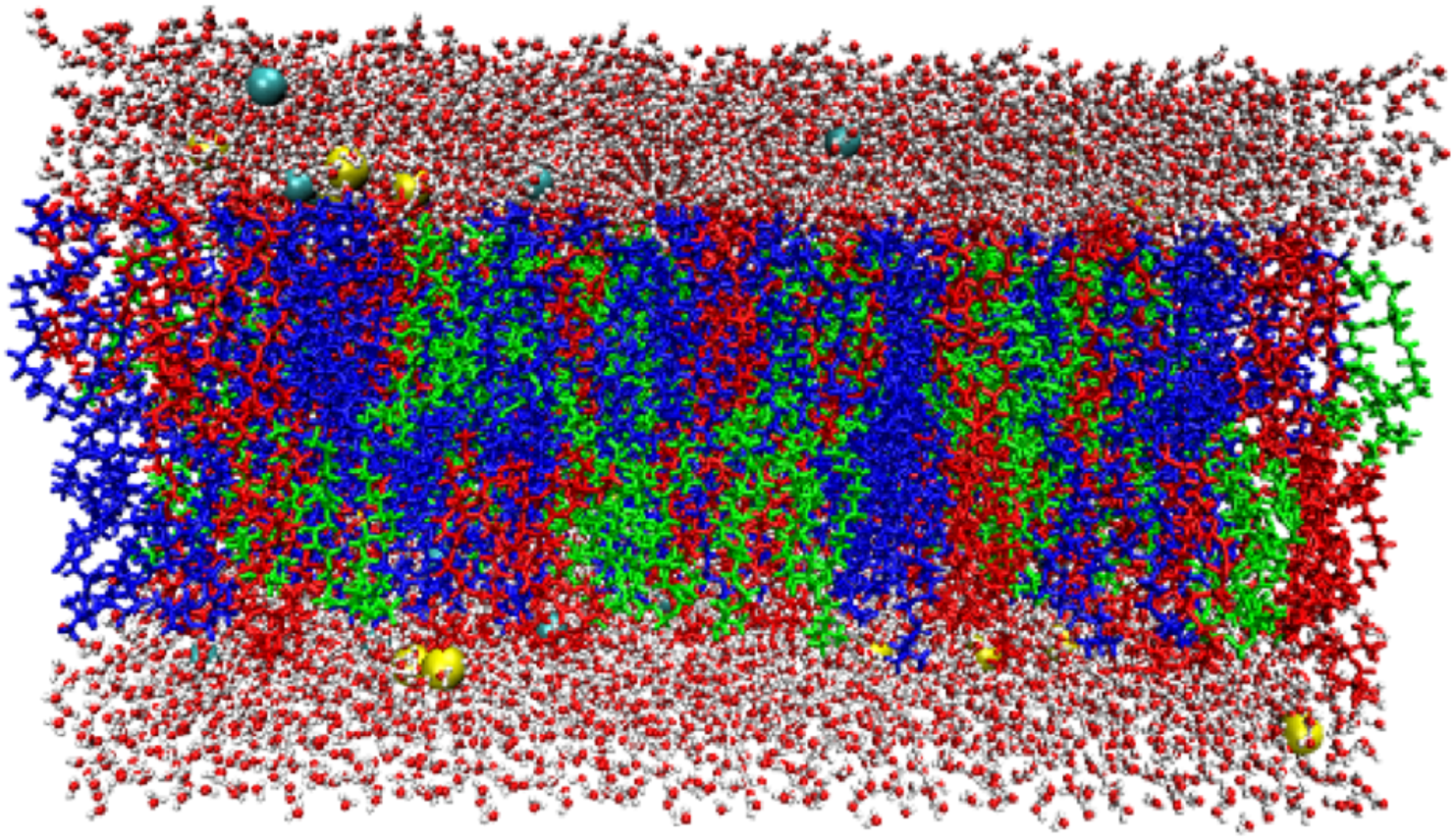
Original lipid membrane constructed by charm-gui membrane builder. The lipids are shown in licorice representation. POPC lipids are shown in green, PLPC lipids are shown in blue and SAPC lipids are shown in red. The water molecules are shown in red-white and the ions are shown as spheres colored in cyan and yellow.

The bilayers were hydrated using 35 water molecules per lipid up to a total of 18,200 water molecules. The ion concentration added to both systems was 0.15 M NaCl. Equilibration was performed initially for 225 ps using the standard six-step CHARMM-GUI protocol (Jo, Kim et al. 2007) with the CHARMM36 lipid force field (Klauda, Venable et al. 2010) and the TIP3P water model (Jorgensen, Chandrasekhar et al. 1983), and was followed by 5 ns equilibration using NAMD (Phillips, Braun et al. 2005).

After bilayer equilibration, a production run was performed in the NPT ensemble for 700 ns using NAMD (Phillips, Braun et al. 2005). The time step was set at 2 fs using the SHAKE algorithm. The van der Waals interactions were smoothly switched off at 12 Å by a force-switching function, and long-range electrostatic interactions were calculated using the particle-mesh Ewald method. The MD simulations were carried out at a constant pressure (1 atm) and temperature at 25°C. Here, it should be noted that the phase conversion temperatures of lipid bilayers from gel to liquid are much lower than the maintained temperature during the simulation (https://avantilipids.com/), and thus lipid bilayers simulated here are in the liquid phase. More precisely the phase transition temperature for the POPC is −2°C, for the PLPC −20°C and finally for the SAPC −40°C. NAMD provides constant pressure simulation using a modified Nosé-Hoover method in which Langevin dynamics is used to control fluctuations in the barostat as described in Martyna et al. (Martyna, Tobias et al. 1994), with piston fluctuation control implemented using Langevin dynamics (Feller, Zhang et al. 1995). A constant pressure of P = 1 atm was imposed with barostat oscillation of 50 ps with a barostat damping time scale of 25 fs. Constant temperature conditions were maintained by a Langevin piston method wih a damping coefficient of 1 ps^-1^, applied to non-hydrogen atoms.

The analysis of the trajectory was carried out using MEMBPLUGIN (Guixa-Gonzalez, Rodriguez-Espigares et al. 2014) within VMD (Humphrey, Dalke et al. 1996) to calculate several biophysical properties such as the bilayer thickness, the surface area per lipid, and the deuterium order parameters, *SCD.* The average electrostatic potential was calculated by using the particle mesh Ewald (PME) plugin of VMD. The PME method permits the efficient computation of long-range electrostatic interactions evaluating the reciprocal sum of the smooth particle-mesh Ewald method resulting in a smoothed electrostatic potential. The algorithm approximates point charges using Gaussians with sharpness controlled by the Ewald factor (Aksimentiev and Schulten 2005). The physicochemical properties studied herein were chosen in order to perform comparisons with the liposome experiments. All errors reported here represent standard deviations calculated by dividing the trajectories into 500-ps pieces and calculating the corresponding properties.

## Results and Discussion

### Molecular Dynamics Simulations

To monitor the bilayer equilibration, we first monitored the area per lipid in all systems (Fig. S3). The area per lipid in each system and for each lipid remained stable after 5 ns, after which the bilayer was considered equilibrated. We thus performed our analyses in the next 700 ns.

Bilayer thickness depends on carbon chain lengths, lipid tilt, and degree of unsaturation of the membrane lipid (Rawicz, Olbrich et al. 2000, Kučerka, Nieh et al. 2011). To calculate the thickness of the bilayer, the distance between two atoms at the ends of the bilayer, usually of the phosphorus atoms, is calculated.

MEMPLUGIN measures the distance between the two density peaks, defined as the first and second central moments of the mass density profile of a particular atom (e.g., phosphorus). The average thickness of the bilayer in both systems is 39.5 Å ±1.6 and 39.6 Å ± 2.7 (Fig. S4).

Although there is no identical study to exactly compare this result with previous ones, several simulations have been made with other lipids having some common elements (fatty acids) with the lipids of the present study. Lipids containing palmitic (C16:0) and stearic acid (C18:0) have been shown to result in a 38 Å membrane thickness when calculating the spacing between phosphorus atoms (Venable, Sodt et al. 2014). Others show that on average the thickness of the bilayer containing POPC is 30 Å (Kučerka, Nieh et al. 2011) and 38 Å (Janosi and Gorfe 2010). As we can see in Fig S4, we do not notice significant differences between the two systems. Generally, the thickness of the membrane is also associated with variations in the area per lipid (area occupied per lipid). The larger the area per lipid the thinner the thickness of the bilayer (Reddy, Warshaviak et al. 2012). However, the correlation between thickness and area per lipid is not very clear when carbon chain lengths vary in lipids such as in the present study.

The area per lipid is a fundamental factor in the description of cell membranes. It is an aggregation property that contains information about the phase, fluidity, and degree of condensation. It is also one of the few parameters that can be compared with experimental measurements. Both the total lipid surface and the lipid surface of each lipid species separately were calculated here. Before performing this calculation a key atom, or a triad of atoms, is selected. The x, y coordinates of the previous set of points are projected to a level bounded by the simulation box, which in turn is divided into polygons using Voronoi, using the qvoronoi program from the Qhull package (Barber, Dobkin et al. 1996). In this way, the lipid surface of each polygon is calculated. The area per lipid, as well as the thickness of the bilayer, can also provide information about the equilibration of the membrane; when the area per lipid reaches a plateau value the membrane can be considered equilibrated (Cournia, Ullmann et al. 2007, Porasso and López Cascales 2012).

The lipid surface of System 1 has an average value of 66 Å^2^ ± 0.7 (Fig. 4), which is consistent with other membrane studies of similar composition. The area per lipid for POPC pure membranes ranges between 64 and 68 Å^2^ (Kučerka, Tristram-Nagle et al. 2006, Klauda, Venable et al. 2010). In the case of PLPC, the value is 69 Å^2^, though lipids were simulated in higher temperature (Chaban 2014). The addition of polyunsaturated lipids has been shown to result in higher values of the area per lipid (Stillwell and Wassall 2003). Moreover, Hyvonen et al., studied double layers of four lipid types separately (Hyvönen and Kovanen 2005), and calculated the area per lipid for POPC, PLPC, and PAPC (1-palmitoyl-2-arachidonoyl-sn-glycero-3-phosphocholine), phospholipids, to be 62 Å^2^, 64.8 Å^2^ and 62.4 Å^2^, respectively. SAPC was also used in this paper, exhibiting area per lipid values ranging from 65.3 to 62.3 Å^2^. Here, it should be noted that the results of the above study were carried out using united atoms and bilayers consisted exclusively of one lipid type.

**Fig. 4.**
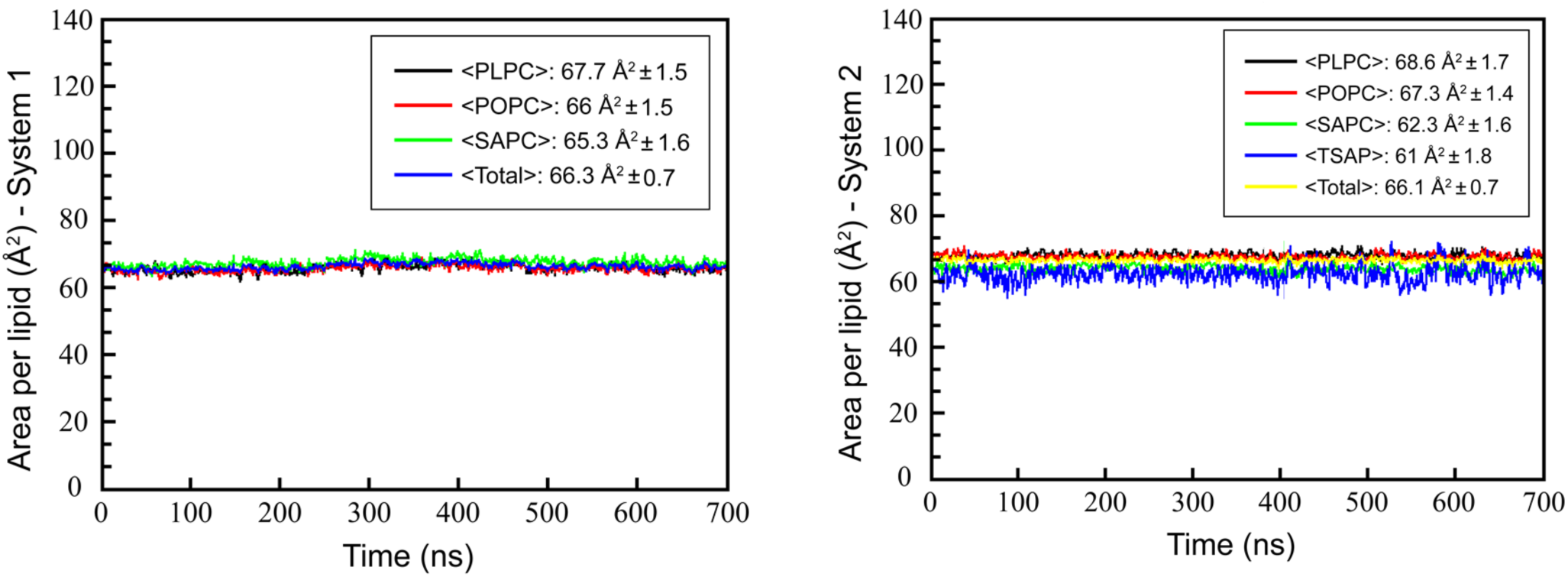
Area per lipid for the two simulated systems. Left panel shows the original constructed System 1 and right panel show the area per lipid of System 2, containing 3% mol. of TSAP.

While the average area per lipid in System 1 and System 2 is identical with values of 66.3 ± 0.7 Å^2^ and 66.1 ± 0.7 Å^2^, respectively, the different lipids exhibit in some cases differences. TSAP has a smaller area per lipid compared to SAPC (61 ± 1.8 Å^2^ vs 62.3 ± 1.6 Å^2^ (Fig. 4)). This smaller area per lipid is consistent with the hypothesis that the *trans* configuration mimics saturated fats, which have a smaller area per lipid compared to unsaturated ones, because the different stereochemistry results to more linear lipid chains. Unsaturated fatty acids are thought to generally increase the fluidity of the membrane and are more agile. In contrast, *trans* fatty acids more resemble saturated ones, which make the membrane more structured and less agile (Alberts, Johnson et al. 2002). Moreover, SAPC assumes a smaller area per lipid in the simulation in System 2, containing the 5-*trans* arachidonic acid (62.3 ± 1.6 Å^2^) compared to the only *cis-*containing phospholipids System 1 (65.3 ± 1.6 Å^2^). This reduction in area per lipid for the SAPC lipid could be attributed to the fact that it associates with the *trans-containing* TSAP lipid and decreases its area per lipid. Smaller areas per lipid can be associated with increased order of the hydrocarbon chains (Cournia, Allen et al. 2015, Angelikopoulos, Sarkisov et al. 2017).

In fact, the structure of lipid hydrophobic tails is directly related to various parameters: condensation of the bilayer and its fluidity as well as its thickness (Lodish and F. 2013). One way to evaluate the ordering of the hydrocarbon tails is to evaluate their order through the deuterium order parameter, -*SCD* (Cournia, Ullmann et al. 2007, Zervou, Cournia et al. 2014, Wang, Gkeka et al. 2016). It is quite important that this parameter can also be measured experimentally by NMR experiments (Petrache, Tu et al. 1999). The *SCD* quantifies the configuration of the hydrophobic tails of the phospholipids by calculating the average of the angles θ for each C-H bond along the bilayer axis-z. This calculation is done separately for each methyl group in each lipid chain by selecting a particular lipid type. The -*SCD* is calculated for each methyl group using the equation:

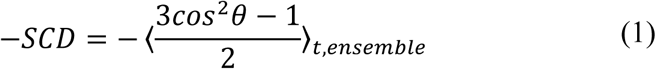

Membranes that are highly ordered exhibit high values of the -*SCD* parameter and vice versa (Vermeer, de Groot et al. 2007). Higher -*SCD* values are present near the polar head of the phospholipids. By contrast, toward the end of the carbon chain lipids are much less ordered.

As mentioned earlier, hydrocarbon chains have higher values near the polar head (carbon C2), while values are much smaller toward the end of the chain (C16, C18, C20). In our study, the *sn*1 chains all contain saturated fats and we do not observe any differences or fluctuations in Fig. 5. In contrast, the *sn*2 chains contain unsaturated fat, and we observe that the -*SCD* values decrease at the double bond (*cis*) location, resulting in a less ordered lipid. For example, the four double bonds of SAPC (C5, C8, C11, C14) show reduced -*SCD* values compared to the *trans* double bond of TSAP at C5, which has an elevated –*SCD* (Fig. 5). This increase in –*SCD* is attributed to the fact that the chain becomes more ordered due to the *trans* geometry, consistent with both experimental data and other MD simulations (Hyvönen and Kovanen 2005). *Trans* fatty acids resemble the structure of saturated fatty acids, which are well-ordered and give higher -*SCD* values, in agreement with the present study. As a result, chains with *trans* geometries are more straight than chains with *cis* geometries and lead to more condensed membrane formation (Lichtenstein 2000) and changes in membrane fluidity (Roach, Feller et al. 2004).

**Fig. 5.**
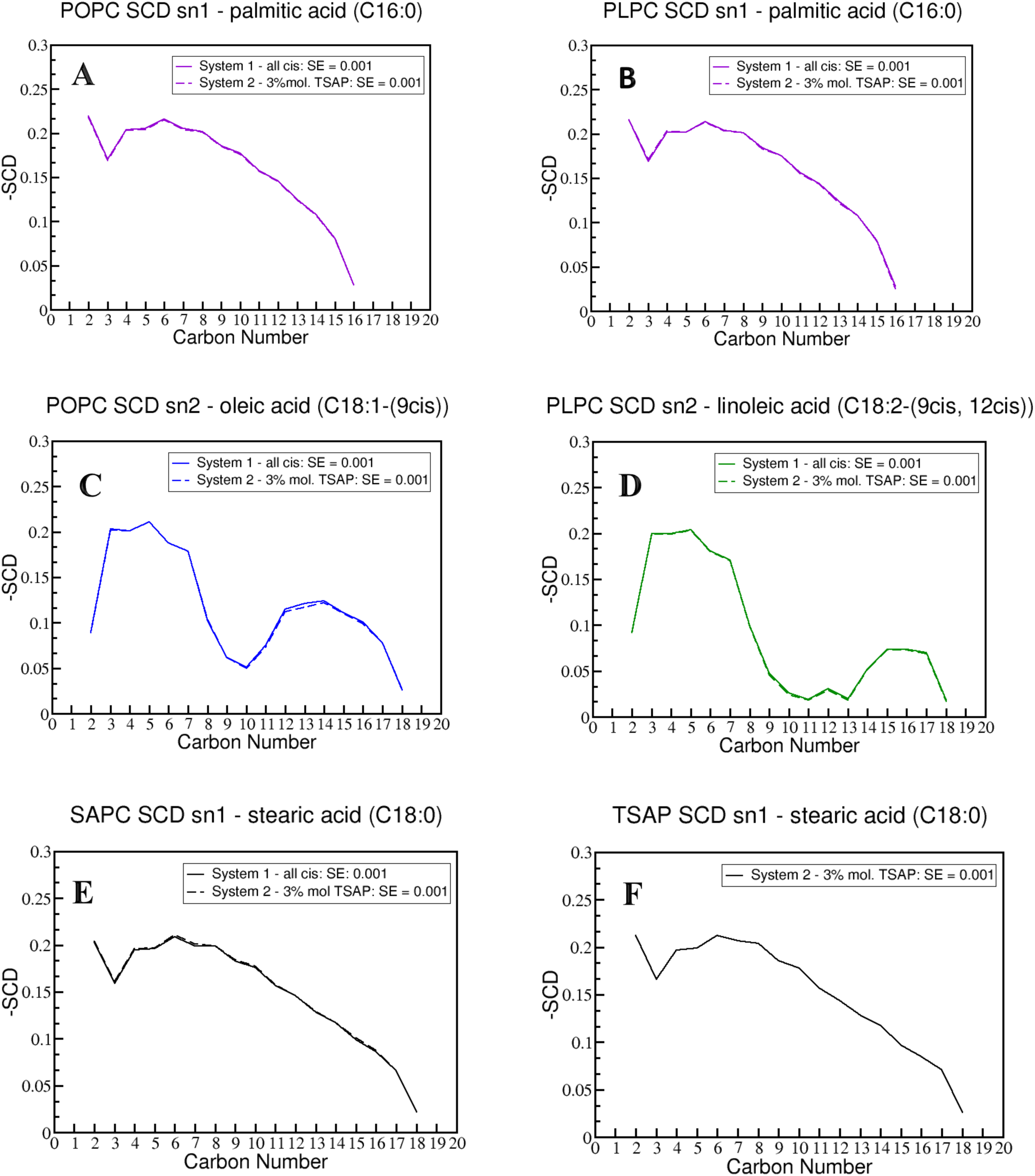
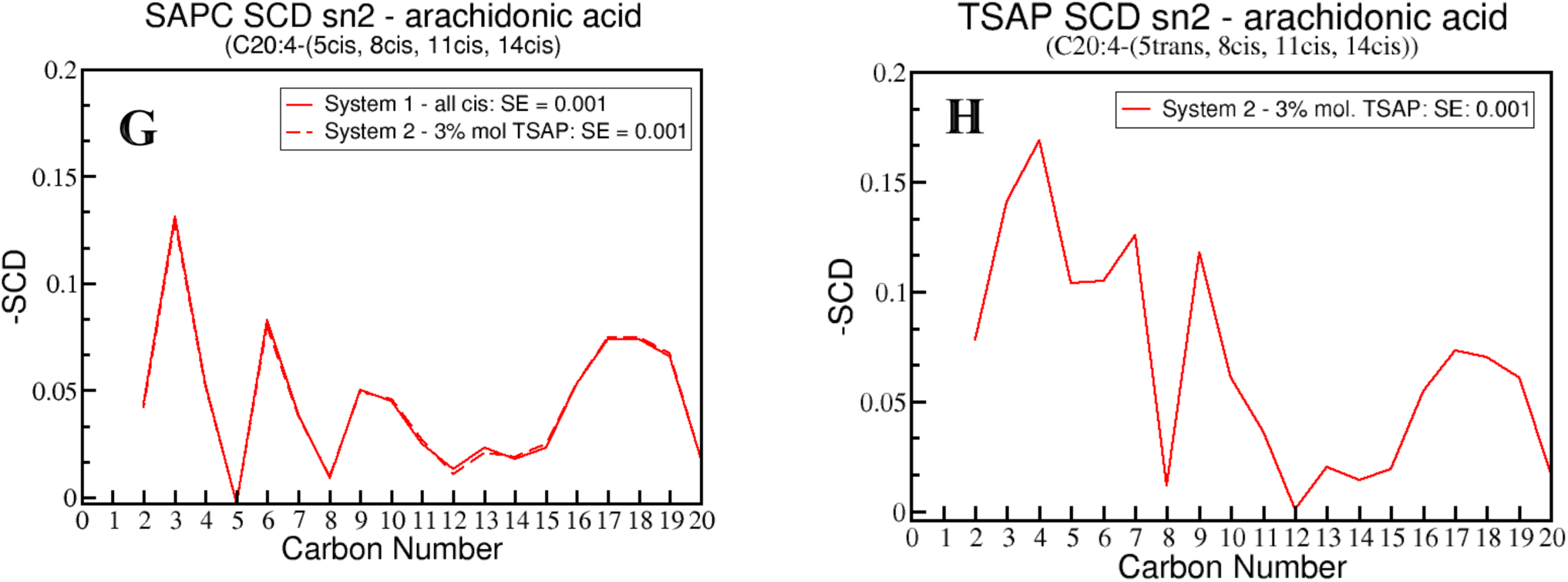
SCD order parameters for A. POPC *sn-*1 chain, B. PLPC *sn-*1 chain, C. POPC *sn-*2 chain, D. PLPC *sn-*2 chain, E. SAPC *sn-*1 chain, F. TAPC *sn-*1 chain, G. SAPC *sn-*2 chain, H. TSAP *sn-*2 chain. A-F monitor both *sn* chains in both simulated systems, while G and H show TSAP only in System 2.

### Liposome Characterization

In order to model the outer membrane of the erythrocytes, the following fatty acids were chosen for our study: palmitic, stearic, oleic, linoleic, and arachidonic. The phospholipids deriving from these fatty acids and used in our study were POPC, PLPC and SAPC (Fig. 2B). Here it should be noted that considering the previous studies in the composition of the lipids our purpose was to make a simple model close to the lipid profile of the fatty acids and study the effect of the cis-trans isomerization in this model.

From a chemical point of view, naturally occurring *cis* isomers can be converted to *trans* isomers by free radicals like RS^•^ radicals (Chatgilialoglu, Ferreri et al. 2014). Fig 1C shows the *cis*-*trans* isomerization process of a double bond catalyzed by thiyl radicals (Chatgilialoglu, Ferreri et al. 2000). Amphiphilic thiols such as HOCH_2_CH_2_SH have been used previously to cause radical stress in biomimetic and cellular models such as liposomes (Chatgilialoglu and Ferreri 2005, Chatgilialoglu and Ferreri 2010). Lipid vesicles such as liposomes have been extensively used in radical-catalyzed *cis*-*trans* isomerization because they are considered representative models for membrane lipid assembly ((New 1990, Chatgilialoglu, Ferreri et al. 2014, Chatgilialoglu, Ferreri et al. 2014). Studies of lipid vesicles have indicated that during radical-catalyzed isomerization there is a regioselectivity on the double bonds closer to the surface favoring the isomers in position 5 and 8 (Ferreri, Samadi et al. 2004). In order to directly compare our simulation and experimental results (restricted to the 5-*trans* isomer of arachidonic acid), we investigate the reaction of HOCH_2_CH_2_S• with SAPC in multilamellar vesicles (MLVs) and the reaction progress was monitored by extraction and transesterification of lipids in order to acquire preferably mostly the 5-*trans* isomer.

Various analytical techniques can be applied to characterize the liposomal structure such as Atomic Force Microscopy (AFM) (Ruozi, Tosi et al. 2007), Dynamic Light Scattering (DLS) (Matsuzaki, Murase et al. 2000), Scanning/Transmission Electron Microscopy (SEM/TEM) (Ruozi, Belletti et al. 2011). AFM belongs to the family of scanning probe microscopy and its resolution approaches 1 Å. The AFM has the ability to explore samples in different environmental conditions, including biological specimens in an aqueous solution such as liposomes. The function of the AFM is based on a tip (probe), which is attached in a flexible cantilever of specific spring constant. The tip is scanning the surface of the sample by contact or non-contact mode depending on the sample. The AFM can provide information on the size and surface properties of the liposomes (Spyratou, Mourelatou et al. 2009).

The morphology and size of the liposomes were observed first with AFM to monitor any differences between System 1 (only *cis*-containing phospholipids) and System 2 (*cis*-and *trans*-containing phospholipids). Liposomes were characterized both with contact and tapping mode on the AFM. When liposomes are placed on mica substrate they are affected and their shape can be altered (Ruozi, Tosi et al. 2007). Another factor that can affect liposomes is the magnitude of the force exerted on them (Bar-Ziv, Moses et al. 1998). Therefore, this can also provide us with information about their elastic properties of the liposomes (Spyratou, Mourelatou et al. 2009). Originally, both liposome systems were observed on a phosphate buffered saline (PBS) buffer solution. However, the liposomes could not be studied due to the effect of PBS on the images. The PBS solution resulted on tree structures that were overlapping the liposomes, making it impossible to see them (Fig. S5), possibly due to salt crystals (Zhu, Sabharwal et al. 2009), which are portrayed through the AFM and overlapping the liposomes. For that reason the liposomes were subsequently studied using pure water.

At first, the morphology of the liposomes of both systems was studied. System 1 produced better topographic images with minimal artifacts and more details in morphology when we operating in the contact mode, while system 2 when we were operating in the tapping mode. This difference may have arisen due to the different stiffness of each system in combination with the spring constant *K* of the AFM cantilever. However, in order to directly compare the two systems, all the dimension measurements were performed using the same mode (tapping mode).

Fig. 6 (a) – (c) shows the topographic images of the liposomes of System 1 (with only cis-containing phospholipids), indicating that liposomes are mainly convex. However, strong bonding forces appear between the liposomes and the AFM tip leading to hysteresis and deformities in the shape of the liposomes indicated by the circles.

**Fig. 6.**
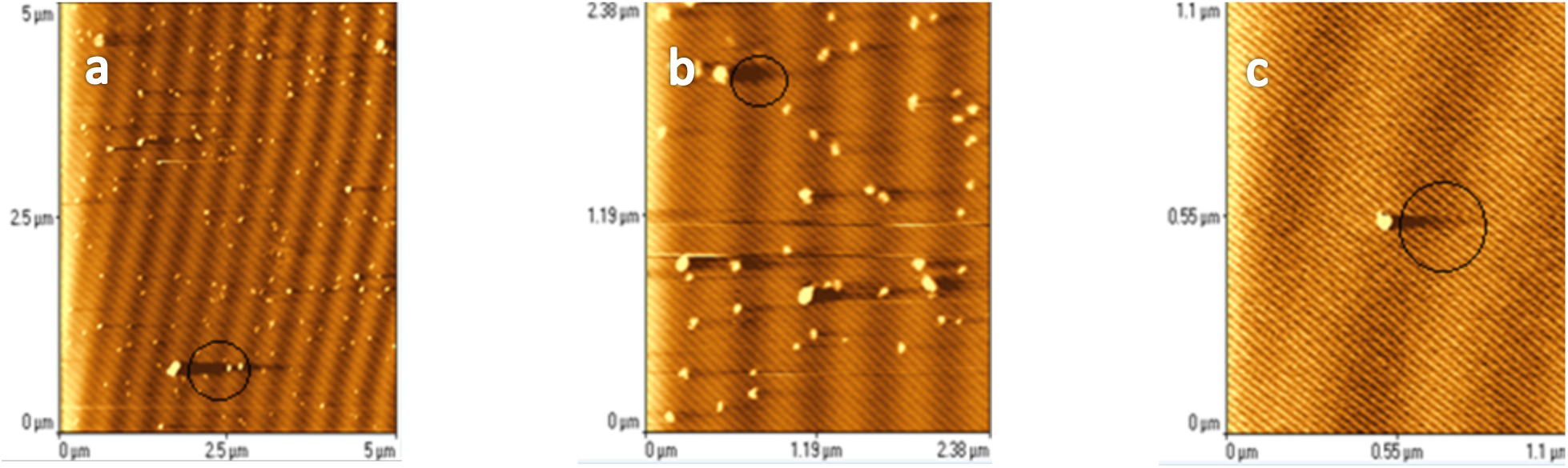
Topographic images of liposomes by contact mode (System 1). The image shows gradual magnification of the circulated area.

Fig. 7 shows the corresponding depiction of liposomes using the lateral forces of the static mode (Lateral Force Microscopy (LFM)), that is, frictional forces generated parallel to the plane of the surface. The LFM provides useful information for the topography in the periphery of the structures. In addition to curved liposomal structures, we also observe hollow-shaped structures with a pressed center and an elevated periphery.

**Fig. 7.**
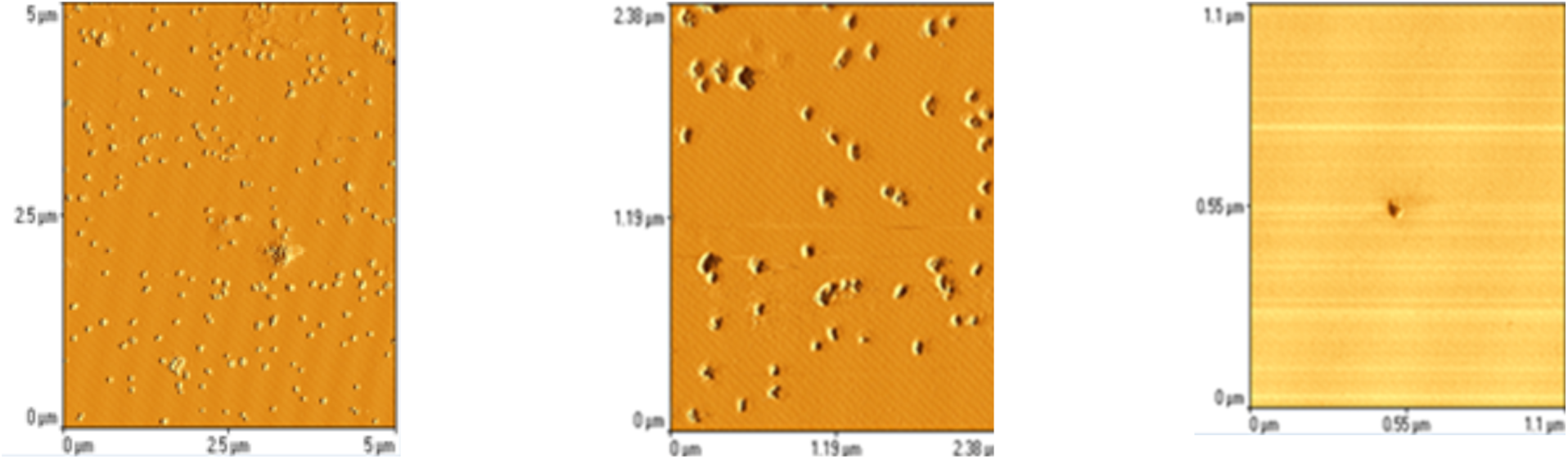
Corresponding topographic images LTM of liposomes (System 1). The images show gradual magnification of the presented area.

For System 2 (with *cis*-and *trans*-containing phospholipids), the images were acquired using tapping mode and the phase imaging method (Fig.8 (a) – (c)). This method can map the composition of the surface according to the local mechanical and structural variations of the sample. We observe that these liposomes are mainly convex, while hollow-shaped structures with compressed center and elevated periphery, as indicated by the arrow in Fig. 8c (donut shape, pancake-like), are also observed. Similar shapes of liposomes have been reported also in other studies (Liang, Mao et al. 2004, Ruozi, Tosi et al. 2007). Also, some liposomes are located in deeper layers in the mica, which indicates absorption by mica. In conclusion, both System 1 and 2 exhibit similar morphologies.

**Fig. 8.**
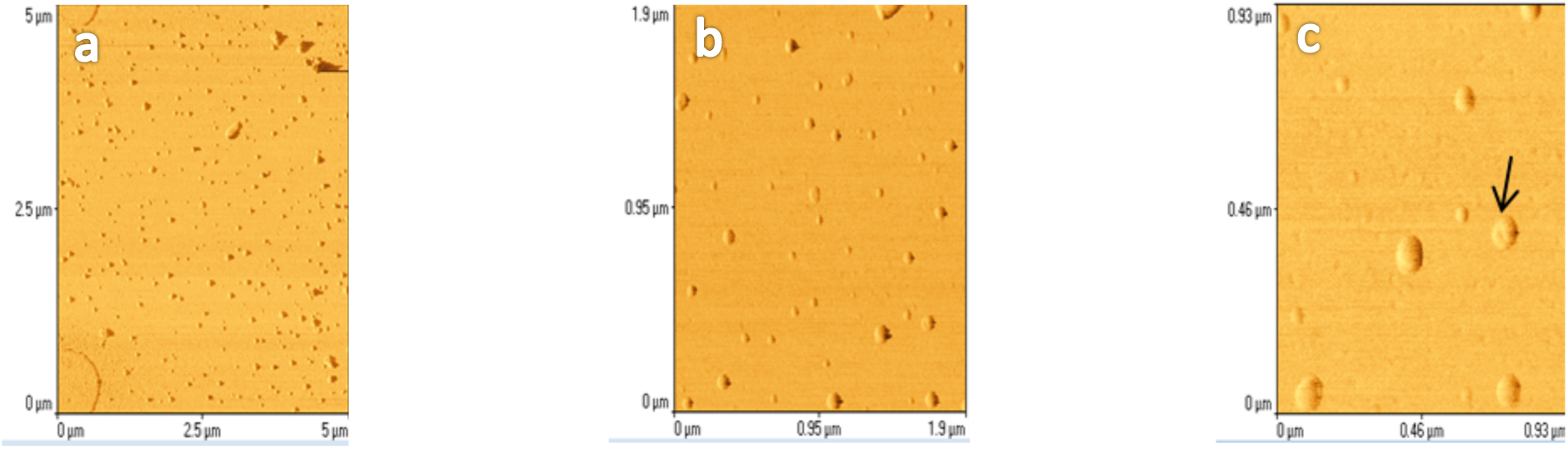
**(a) – (c).** Topographic images of liposomes (System 2). The images were taken using the phase imaging method and they are presented with gradual magnification. The arrow shows the donut –like shape of the liposome.

Subsequently, the diameter of the liposomes was measured to evaluate an average size. The comparative measurement of the diameter of the two types of liposomes was performed using the AFM tapping mode and the dimensions of the liposomes were measured from the topographic images using the AFM SpmLabAnalysis software. The diameter of the liposomes is calculated from the spatial width of the peak from its base according to Fig. S6. The measurement of the width of the diameters (green arrows) and the height (blue) of the liposomes using the SpmLabAnalysis software can be seen in the supplementary material. The average size diameter of the liposomes of System 1 was measured to be 120nm ± 20 nm, while the average diameter of System 2 was measured to be 100 ± 15 nm (see Fig. S7).

Comparatively, *trans*-containing liposomes (System 2) appear to be smaller in size than only *cis*-containing liposomes (System 1) That is consistent with the observation that *trans* lipids are more ordered and have a smaller area per lipid, thus resulting to a more condensed liposome. Moreover, AFM spectroscopy measurements were performed to compare the elasticity of the two systems. Force-distance curve were produced by recording the liposome pushback on the tip (deflection) versus vertical z position of the tip probe. The difference in the slope of the force curves represents the difference in stiffness of the two systems. The slope of the linear curve, which fits the force versus indentation plot, was used to calculate the Young’s elastic modulus, E, according to Hertz-Sneddon model (Sneddon 1965, Cappella and Dietler 1999):

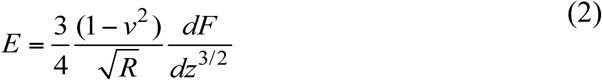

where, *v* is the Poisson ratio, *R* the radius of the AFM tip, *F* the loading force and *dz* the indentation of the AFM tip into the liposome. A tip radius of 40 nm diameter and cantilever spring constant of 0.02 N/m was used for all calculations, and the typical radius of the tips was specified by the manufacturer. Lipid bilayer membranes are generally assumed to be quasi incompressible, characterized by a Poisson ratio of 0.50. Figure S7 illustrates the force curve as a function of the tip indentation for the two systems. The force versus indentation^3/2^ graph follows a linear trend (Fig. S8) and the slope was fitted as *dF/dz*^*3/2*^ to approximate the Young moduli. The Young moduli were measured to be 1.13 ± 0.01 MPa and 2.27 ± 0.01 MPa for System 1 and System 2, respectively. Based on these measurements *trans*-containing liposomes appear less elastic than *cis*. At least two or three measurements were performed on both samples with contact and tapping mode and many different locations of the samples were scanned by AFM each time.

The dimensions of the liposomes calculated using the AFM are consistent with those measured by the standard Dynamic Light Scattering (DLS), which follows next. DLS is a non-invasive technique for measuring the size, the poly dispersion index and the zeta potential of molecules and particles in a suspension. The average size and size distribution obtained from DLS can easily be combined with AFM measurements.

DLS was used along with AFM to measure the average size and characteristics of the two liposome systems and for acquiring better statistics as DLS measures liposomes in solution. The size of the liposomes in both systems as referred previously was performed after filtration. In Fig. 9 (a) – (b) the size distribution of both Systems 1 and 2, respectively, are shown.

**Fig. 9.**
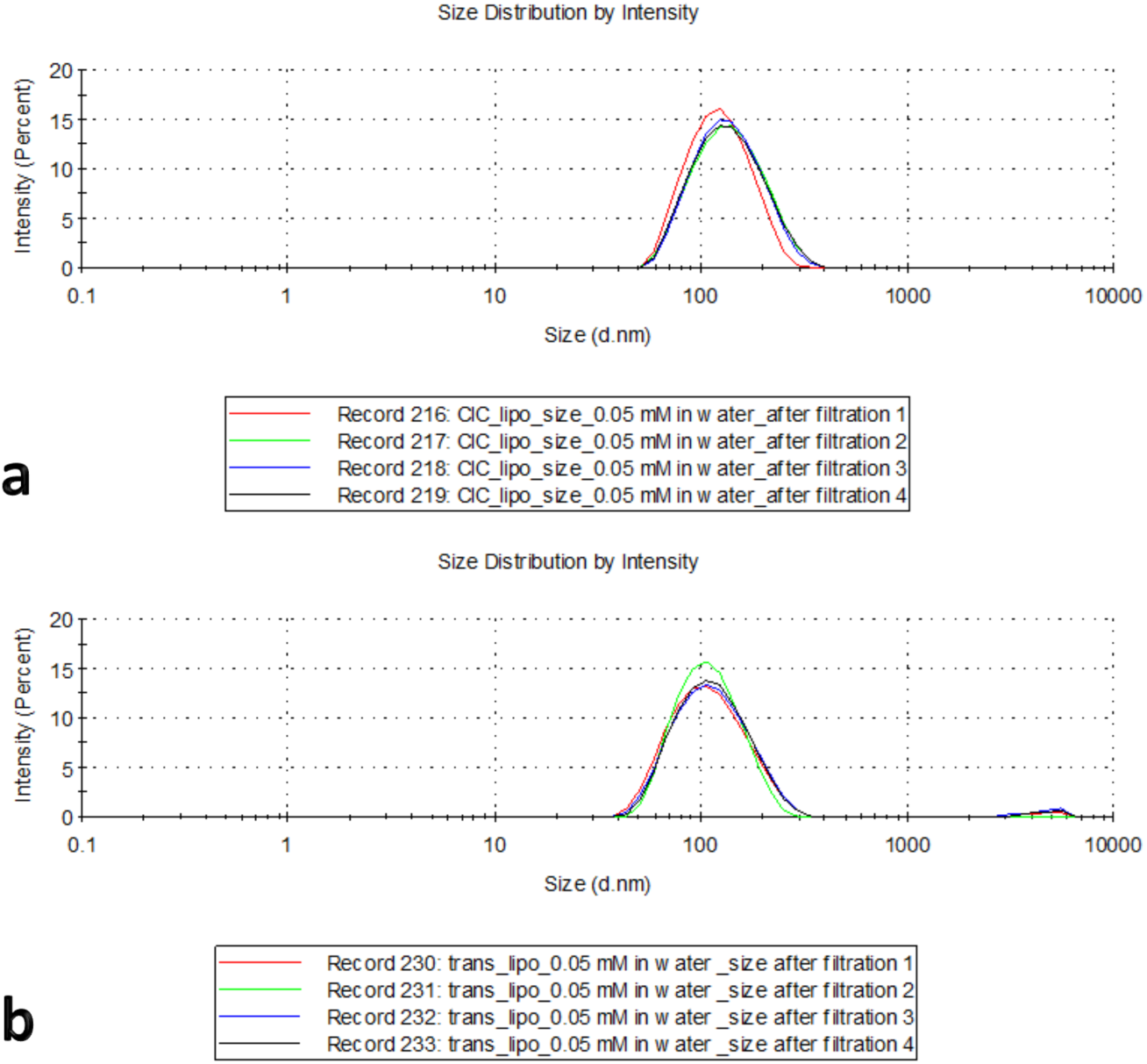
(a) Size Distribution of the liposomes of System 1, (b) Size distribution of the liposomes of System 2.

The average diameter of the synthesized liposomes was 123 ± 6.1 nm for System 1 and 104.6 ± 2.0 nm for System 2. Following the size distribution, the zeta potential was calculated (Fig. 10 (a) – (b)). The zeta potential for System 1 was −26.9 ± 3 mV, with pdi= 0.130 ± 0.015 and for System 2 the zeta potential was −11.2 ± 2.98 mV with pdi= 0.174 ± 0.021.

**Fig. 10.**
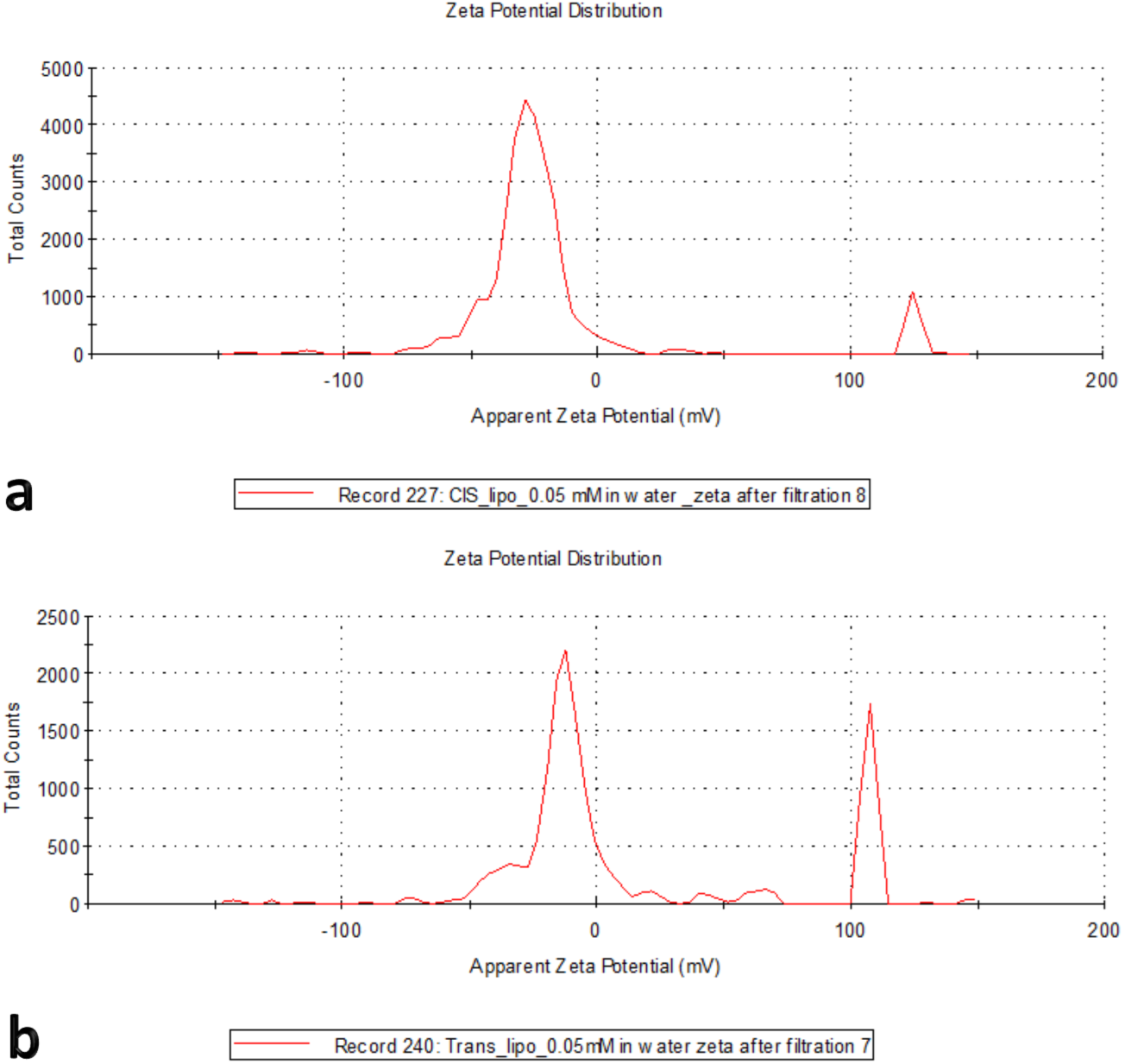
(a) Zeta Potential of System 1, (b) Zeta Potential of System 2.

Calculation of the electrostatic potential of the two Systems with MD simulations shows that the trans-containing arachidonic acid membrane exhibits the lowest potential variation across the membrane, with 2.2 mV, while the all-cis arachidonic acid system varies by 2.6 mV.

From the measurements of both systems we conclude that both solutions were colloid. The average size of System 1 is ∼18 nm bigger than System 2, consistent with the AFM measurement. Again, the difference in size can be explained due to the different configuration on the 5-double bond of the *trans* arachidonic acid. System 2 renders the lipid more well-ordered resulting to tighter packing of the lipids and smaller liposome size. Furthermore, System 2 appears with a more homogeneous distribution as we can see from the diagrams. From the comparison of the two types of liposomes we observe that System 2 exhibits lower potential but improved size probably due to its stereochemistry.

## Conclusions

Membrane properties are driven by the presence of unsaturations, needed for the functioning of channels, receptors and proteins (Ferreri and Chatgilialoglu 2012). Although *cis* fatty acids have been evolutionarily selected by eukaryotes for incorporation into membranes, *trans* fatty acids that are associated with adverse health effects can be also taken in from exogenous sources or may form due to endogenous oxidative stress. Arachidonic acid is an important functional element of cells, and contains a high degree of unsaturation. Because its 5-*trans* geometrical isomer is formed endogenously, we chose it here in order to study the effect of *trans* fatty acids in model membranes. We investigated a model system containing 5-trans arachidonic acid in a realistic concentration and membrane composition, resembling those found in nature. We show that incorporation of *trans* lipids in model membranes affects the physicochemical properties of the membrane (size, ordering, area per lipid), which may in turn affect various functions of the cell.

*Trans* fats are believed to resemble the structure of saturated lipids, as the *trans* geometry forces the lipid to adopt a more ordered orientation in the membrane. Our MD simulations of an all-*cis* model erythrocyte membrane containing POPC, PLPC, and SAPC compared to the same model membrane with incorporated 3% mol. of TSAP, showed that the NMR lipid order parameters, -*SCD*, of the *trans* double bond at position C5 are higher than the *cis* counterpart. This *cis* to *trans* change in geometry causes the TSAP lipids to become more arranged and packed closer to one another, resulting to a significantly decreased area per lipid of TSAP compared to SAPC as observed in the simulations. The measurements for both *SCDs* and area per lipid are consistent with previous lipid studies both in mono and poly unsaturated lipids (Soni, Ward et al. 2009, Kulig, Pasenkiewicz-Gierula et al. 2016), however, this is the first time that a model erythrocyte membrane and the effect of 5-mono *trans* isomer of arachidonic acid on its physicochemical properties is reported. The smaller area per lipid and higher ordering of the membrane observed in the MD simulations are further supported by AFM and DLS measurements for liposome size, which show that *trans*-containing liposomes adopt a smaller size. Using AFM we also studied the morphology of liposomes, which show that, trans-containing liposomes are stiffer than *cis*, which were found to be elastic.

In conclusion, inclusion of *trans* arachidonic acid on a model liposomal membrane in a biologically-relevant concentration, affected the physicochemical properties of the model erythrocyte membrane that we studied here, such as membrane rigidity, area per lipid, order parameters. Studying *trans* fatty acids that are produced with endogenous isomerization processes by free radicals can provide new insights into the effect of *trans* lipids in the human organism. Furthermore, the presence of radical-catalyzed products such as mono-*trans* isomers is intriguing because fatty acid moieties may play a key role as biochemical markers of signaling, degenerative processes or *in vivo* isomerization processes, provided that we can carefully monitor the geometrical and positional isomers (Ferreri, Faraone Mennella et al. 2002). This study is a first step into understanding the mechanism with which *trans* fatty acids regulate the physicochemical properties of membranes, but long-term research is still needed to elucidate the mechanisms with which *trans* fatty acids may be implicated in human diseases.

## Acknowledgements

We acknowledge computational time granted from the Greek Research & Technology Network (GRNET) in the National HPC facility – ARIS under project ID pr001026-MAG-NANO-MEM. We also acknowledge Dr. Eleni Efthimiadou for providing us the Zetasizer for the Dynamic Light Scattering experiments and Giorgia Giacometti for her ancillary help in the liposome preparation. ZC was co-funded by the European Commission under the H2020 Research Infrastructures Contract No. 675121 (project VI-SEEM).

## Compliance with Ethical Standards

Conflict of Interest: All authors declare that they have no conflict of interest.

Ethical approval: This article does not contain any studies with human participants or animals performed by any of the authors.

